# Improved performance of biohybrid muscle-based bio-bots doped with piezoelectric boron nitride nanotubes

**DOI:** 10.1101/2022.03.30.486204

**Authors:** Rafael Mestre, Judith Fuentes, Laura Lefaix, Jiaojiao Wang, Maria Guix, Gonzalo Murillo, Rashid Bashir, Samuel Sánchez

## Abstract

Biohybrid robots, or bio-bots, integrate living and synthetic materials following a synergistic strategy to acquire some of the unique properties of biological organisms, like adaptability or bio-sensing, which are difficult to obtain exclusively using artificial materials. Skeletal muscle is one of the preferred candidates to power bio-bots, enabling a wide variety of movements from walking to swimming. Conductive nanocomposites, like gold nanoparticles or graphene, can provide benefits to muscle cells by improving the scaffolds’ mechanical and conductive properties. Here, we integrate boron nitride nanotubes (BNNTs), with piezoelectric properties, in muscle-based bio-bots and demonstrate an improvement in their force output and motion speed. We provide a full characterization of the BNNTs, and we confirm their piezoelectric behavior with piezometer and dynamometer measurements. *We* hypothesize that the improved performance is a result of an electric field generated by the nanocomposites due to stresses produced by the cells during differentiation, which in turns improves their maturation. We back this hypothesis with finite element simulations supporting that this stress can generate a non-zero electric field within the matrix. With this work, we show that the integration of nanocomposite into muscle-based bio-bots can improve their performance, paving the way towards stronger and faster bio-hybrid robots.

## Introduction

Biological systems have evolved throughout millennia to develop sophisticated mechanisms of self-organization, actuation and sensing that have been difficult to replicate in the field of robotics. In particular, biomimetic soft robotics has emerged as a promising sub-field of the robotics that aims at overcoming some of the challenges related to the mechanical properties of their materials, such as compliance, flexibility and overall safety for human interaction. Commonly, the rigidity and stiffness of conventional materials in robotics limit their applications into certain healthcare or biomedical disciplines (Patino, Mestre, and Sánchez 2016; Cianchetti et al. 2018; Ricotti and Menciassi 2012). Recent developments in material science have made possible the fabrication of biomimetic soft robots that are able to perform some simple types of actuation, (Wehner et al. 2016) including crawling (Shepherd et al. 2011; Tolley et al. 2014) or grasping (Martinez et al. 2013), but they are still far from the degree of complexity and movement sophistication in living organisms.

One of the most investigated applications of soft robotics is the development of artificial muscles that can mimic the performance of native muscle tissue in mammals. Muscle tissue is inherently complex, being simultaneously strong and fast while enabling a wide variety of movements through an efficient self-organization of its fiber bundles. However, current materials still lack the ability to fully replicate these properties (Ilievski et al. 2011). Even more, other features from biological tissues, such as self-healing, energy efficiency, power-to-weight ratio, adaptability or bio-sensing, to name only a few, are strongly desired but difficult to achieve with artificial soft materials (Mestre, Patiño, and Sánchez 2021). Bio-hybrid robotics is born at this point as a synergistic strategy to integrate the best characteristics of biological entities and artificial materials into more efficient and complex systems, hoping to overcome the difficulties that current soft robots face. Several strategies to unify the development of bio-hybrid devices have emerged, such as the robotic taxonomic key (RTK), that identifies four main elements that an autonomous robotic system can have: structure, actuation, sensing and control (Webster-Wood et al. 2017). From this starting point, one can analyze the capabilities of a bio-hybrid system, defining the artificial or organic nature of the four key elements, to help in the classification and further development on the field. By finding more efficient ways of unifying the most suitable features of current technologies and living entities, the field could reach a change of paradigm to boost the performance and efficiency of robotic systems.

Initial examples of bio-hybrid robots based on skeletal muscle tissue were bio-actuators based on the deflection of cantilevers by single myotubes or full tissue (Sakar et al. 2012; Das, Wilson, and Hickman 2007; Shimizu et al. 2010; Boudou et al. 2012; Hoshino and Morishima 2010), or used as grippers (Kabumoto et al. 2013; Morimoto, Onoe, and Takeuchi 2018; 2020). Untethered bio-hybrid robots, on the other hand, have been mainly based on crawling mechanisms (Raman and Bashir 2017), although some others have relied on swimming (Guix et al. 2021; S. J. Park et al. 2016). Cvetkovic *et al*. developed a 3D-printed bio-bot composed of two legs joined by a beam structure that could walk on the bottom of a Petri dish by differences in friction between two asymmetric legs (Cvetkovic et al. 2014). Indeed, they demonstrated, both experimentally and by simulations, that adding an asymmetry in the design was necessary for motion. Later on, Raman *et al*. presented the same bio-robotic device controlled by light after optogenetically modifying the skeletal muscle cells to contract upon blue light illumination, obtaining a cable-free and remote control system (Raman et al. 2016). In addition to the unidirectional locomotion, Wang *et al*. designed and fabricated a dual-muscle-ring biohybrid walker through a systematic approach based on modeling and simulation. Maneuverability was indeed demonstrated through incorporating two independent muscle actuators on a computationally selected 4-legged scaffold (Wang et al. 2021). Other examples with the same platform have been dedicated to the investigation of some of the features that bio-hybrid integration can offer, such as self-healing (Raman and Bashir 2017), adaptability (Raman et al. 2016), the addition of neurons (Cvetkovic, Rich, et al. 2017), their long-time preservation (Cvetkovic, Ferrall-Fairbanks, et al. 2017; Grant et al. 2019), their scalability (Pagan-Diaz et al. 2018) or their integration with micro-electrodes (Y. Kim et al. 2020). In particular, recent efforts are being dedicated to the integration of neuronal tissue and skeletal muscle tissue, in order to better resemble native muscle and obtain an improved controllability of these bio-robots (Aydin et al. 2020; Kaufman et al. 2020). On this subject, recently, Aydin *et al*. presented a bio-hybrid swimmer with functional NMJ that could swim with flagellar dynamics, although at low speeds due to the difficulty of motion at low Reynolds number (Aydin et al. 2019).

Advances in the field of smart materials could bring another level of complexity and applicability into this kind of devices with the integration of responsive or nanocomposite hybrid substrates (Sydney Gladman et al. 2016; Khoo et al. 2015; Gaharwar, Peppas, and Khademhosseini 2014). Gold nanoparticles or nanowires, for instance, have been widely investigated for the development of cardiac patches (Dvir et al. 2012; Zhu et al. 2017), but little attention has been paid to their applications in skeletal muscle tissue bioengineering or hybrid bio-robots based on them (Marino et al. 2017). Likewise, graphene, graphene oxide (GO) or reduced graphene oxide (rGO) can provide benefits in tissue engineering of skeletal muscle thanks to improvements in the mechanical and conductive properties of the scaffolds (Ahadian et al. 2014; Palmieri et al. 2020). For instance, Du *et al*. used rGO in a biodegradable polycitrate-based elastomer, increasing its tensile strength and improving the differentiation and regeneration of skeletal muscle cells through the physicochemical properties or rGO (Du et al. 2018). GO has increased the viability and metabolic activity of encapsulated C2C12 cells in GO-alginate scaffolds (Ciriza et al. 2015) and graphene sheets have been used to regulate the differentiation of skeletal muscle cells in 2D (Ahadian et al. 2014) or to control the actuation of bio-bots as embedded electrodes (Y. Kim et al. 2020). Recently, gold nanoparticles embedded in hyaluronic acid scaffolds have also been used to improve the differentiation and thus the motion of bio-hybrid robots (D. Kim et al. 2022).

Piezoelectric nanocomposites, on the other hand, could be excellent candidates for the next generation of nano-engineered bio-hybrid robots, due to the possibility of converting sound waves into voltage differences and providing remote actuation of skeletal muscle tissue (Vargas-Estevez et al. 2017; 2018). Moreover, as electrical stimulation during myogenesis can improve the differentiation and maturation of skeletal muscle (Fujita, Nedachi, and Kanzaki 2007; Mestre, Patiño, Barceló, et al. 2019), the addition of piezoelectric nanocomposites could offer benefits in tissue engineering. The mechanical stress generated by cell-laden hydrogels could be transmitted to the piezoelectric nanocomposites, that in turn generate a piezo potential along the nanostructure, creating a charge redistribution around the cell membranes. The use of nanogenerators (NG), in the way of piezoelectric nanostructures, for the stimulation of bone and muscle cells has been recently reported (Blanquer et al. 2022; Murillo et al. 2017). The electromechanical NG–cell interaction triggered the opening of ion channels present in the plasma membrane of osteoblast-like cells (Saos-2) and muscle cells inducing intracellular calcium transients, which at the same time is known to regulate muscle contraction. Poly(vinylidene fluoride) (PVDF) has also been explored for bone and neural tissue engineering (Li, Liao, and Tjong 2019). For instance, Kitsara *et al*. used poly(vinylidene fluoride) (PVDF) to mimic the inherent piezoelectric properties of bone, also observing an increase in intracellular calcium compared to control samples (Kitsara et al. 2019). Piezoelectric boron nitride nanotubes (BNNTs) (Tiano et al. 2014), although they have received less attention in biomedical applications compared to carbon nanotubes (Pok et al. 2014; Sirivisoot and Harrison 2011; Ahadian et al. 2015), have been used in combination with skeletal muscle cells in 2D cultures to improve the differentiation. Ciofani *et al*. investigated the interactions between BNNTs coated with poly-L-lysine and C2C12 cells, finding an increased protein synthesis and no adverse effects in differentiation markers like MyoD or fusion index of myotubes, demonstrating their suitability for biomedical applications (Ciofani et al. 2010). Later on, Ricotti *et al*. described engineered polyacrylamide gels with micro-grooves in which fibroblasts and myoblasts were co-cultured. They supplemented the culture medium with BNNTs and applied ultrasonic stimulation to the samples, thus creating stress on the nanotubes that would be translated into local electric fields that could stimulate the cells, achieving longer, thicker and more functional myotubes (Ricotti et al. 2013).

These promising results, only obtained in 2D cultures, point towards the benefits of the addition of BNNTs in three-dimensional cell-laden hydrogels. For this reason, in this work, we study the integration of this type of piezoelectric nanocomposites into skeletal muscle tissue constructs to improve the performance and force generation of bio-robots. We demonstrate that the integration of BNNT nanocomposites into bio-bots provides faster walking speeds and stronger force output when compared to control samples, thus supporting the hypothesis that self-stimulation of the myotubes in the form of spontaneous contractions and passive compaction that take place during differentiation can induce piezoelectric effect that translates the mechanical stress from the muscle tissue into electrical stimulation, resulting in an efficient feedback loop stimulation. To support our hypothesis, we characterize the BNNTs to prove their piezoelectric effect and we perform finite element analysis (FEA) simulations to show this effect.

## Results and discussion

### Fabrication of bio-bots

Biological robots are becoming increasingly complex by the integration of different types of cells, like neurons (Cvetkovic, Rich, et al. 2017; Aydin et al. 2020; 2019), the design of more sophisticated structures (S.-J. Park et al. 2016), the accomplishment of controlled and complicated tasks (Morimoto, Onoe, and Takeuchi 2018) or the integration of smart materials and designs or nanocomposites into the skeletons (Y. Kim et al. 2020; Guix et al. 2021). However, some of the methods to obtain three-dimensional skeletal muscle tissue with contractile capabilities are already rather established in the literature. We use the method of Raman et al. (Raman, Cvetkovic, and Bashir 2017). Briefly, a myoblast-laden hydrogel composed of Matrigel, fibrinogen and thrombin, was casted into an injection mold made of PDMS (Figure 1A). Later, after 2 days in GM supplemented with ACA, the tissue was gently transferred into a skeleton made of PEGDA, created *via* digital light processing (DLP) 3D printing. This skeleton was based on previous designs reported initially by Cvetkovic *et al*., consisting on two small T-shaped legs joined together by a thin bridge (Cvetkovic et al. 2014). The mechanism of motion was based on friction between the legs and the substrate (Pagan-Diaz et al. 2018). PEGDA is a porous hydrogel that sinks and therefore the bio-bot can walk on the surface. An asymmetry was introduced in the design by making one of the legs longer, as can be seen in Figure 1A (center). In this way, the difference between friction coefficients on both sides of the bio-robot produced a net movement towards the side of the longer leg (Cvetkovic et al. 2014; Pagan-Diaz et al. 2018). Figure 1B shows several microscope images of an assembled bio-robot facing upwards (left), where the base of its legs can be observed, sideways (center) and in walking position (right).

**Figure 1.**
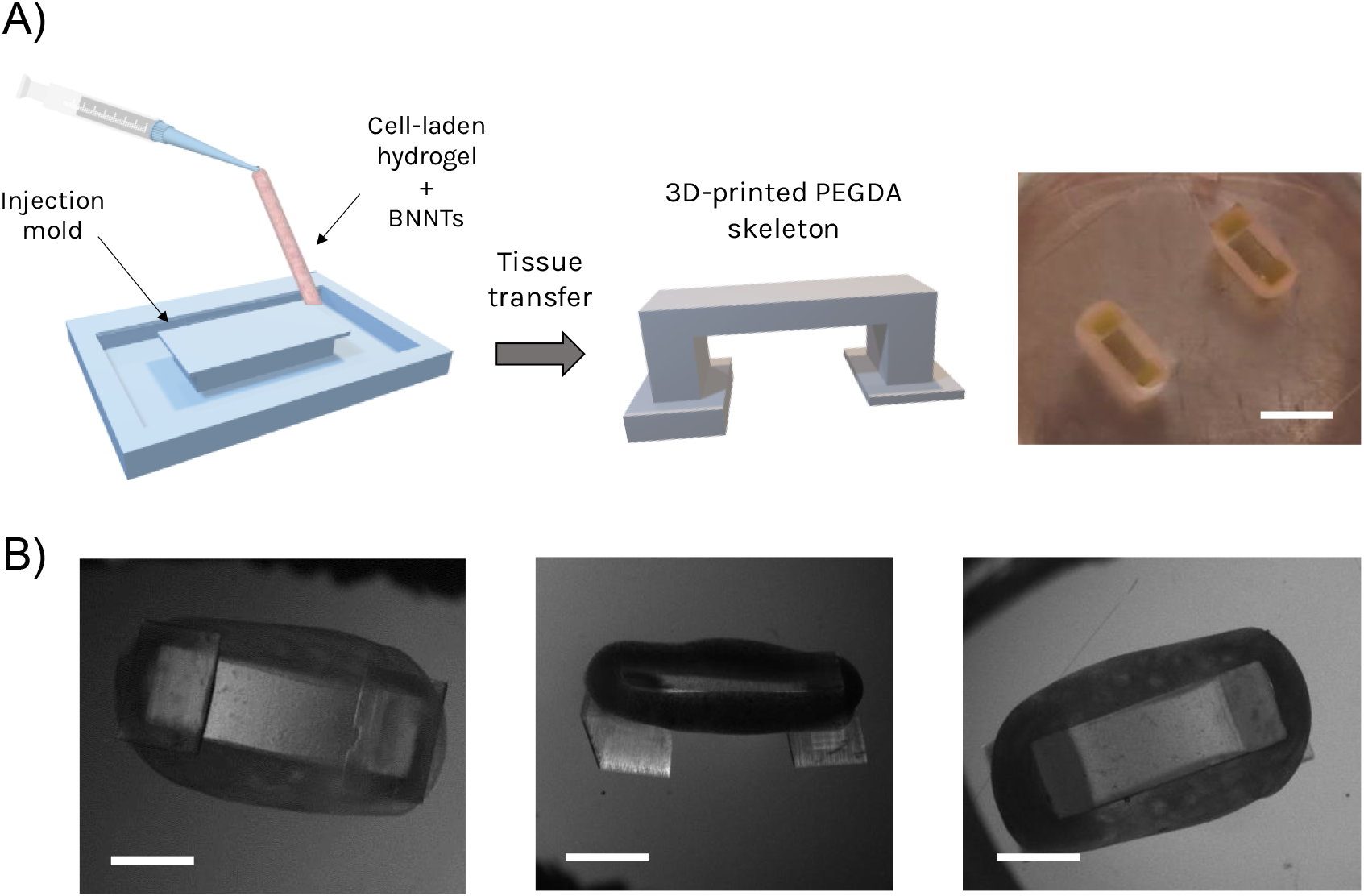
Fabrication of BNNT-loaded bio-bots. A) The cell-laden hydrogel with BNNTs is loaded into an injection mold and left to crosslink for 2 days and it is then transferred into a 3D-printed PEGDA skeleton. Scale bar: 5 mm. Snapshots of a fully formed bio-bot in three different positions: facing up (left), on its side (left), and in walking position (right). Scale bars: 3 mm.

BNNT-loaded bio-bots were fabricated following the same cell encapsulation protocol but dispersing the nanocomposites in the hydrogel before the mold casting process at a concentration of 5 μg/mL (Figure 1). This concentration was chosen since it was previously demonstrated to be biocompatible and show the highest increase in protein content per μg of DNA, when compared to 0 μg/mL and 10 μg/mL (Ciofani et al. 2010). While being kept in ice after sonication, 2.86 μL of a 0.5 mg/mL stock solution of BNNTs in ethanol was mixed with the hydrogel and homogenized thoroughly with the pipette, without creating any bubbles. Then, the BNNT- and cell-laden hydrogel was manually casted into the injection molds and left in a cell incubator for 1 h, until Matrigel was fully crosslinked. Then, GM supplemented with ACA was added and maintained for two days, just like control bio-bots. Then, all tissue constructs were gently lifted from the molds and transferred around the PEGDA skeletons, while their media was changed to DM supplemented with ACA.

The addition of nanocomposites into cell-laden hydrogels have been demonstrated to provide beneficial aspects in the differentiation or maturation of muscle cells. In particular, conductive nano-composites, such as gold nanoparticles (Marino et al. 2017) or nanowires (Dvir et al. 2012), graphene (Ahadian et al. 2014), GO (Ciriza et al. 2015) or rGO (Ku and Park 2013) can improve the mechanical properties of the scaffolds, provide mechanical cues for better alignment, as well as enhance their conductivity for a more efficient propagation of depolarization potentials, which can help in the synchronization of cell contractions (Palmieri et al. 2020). Piezoelectric nanocomposites provide an additional level of benefits that can be considered dynamic due to the combination of piezoelectricity and the active and passive forces exerted by skeletal muscle tissue during myogenesis. The piezoelectric effect is the creation of voltage difference in certain materials when a mechanical strain is applied or, inversely, the deformation of a material when an external electric field is applied. Skeletal muscle tissue can generate two types of stress: a passive stress during myogenesis when myoblasts pull from the surrounding hydrogel chains and compact the tissue, which helps in the self-assembly of bio-actuators or bio-robots; and an active stress in the form of contractions, which can be induced by electric fields but can also occur spontaneously during tissue differentiation (Mestre, Patiño, Barceló, et al. 2019; Cvetkovic et al. 2014). The electric output of the integrated piezoelectric nanotubes within the cell-laden hydrogel depends on the physical parameters of the cell-laden hydrogel and the BNNTs. The piezoelectric coefficient of the BNNT is crucial for the overall piezoelectric behaviour (H. S. Kim et al. 2018). For these reasons, the integration of piezoelectric nanocomposites within the cell-laden hydrogel could potentially transform the stresses generated by the tissue into small voltage differences that could, in turn, provide electrical stimulation to the myotubes during differentiation, in a feedback loop.

### Characterization of BNNTs

BNNTs are a type of one-dimensional structures, similar to carbon nanotubes (CNTs), which have gained significant attraction in the past years due to their outstanding properties. For instance, they possess high chemical stability (Golberg et al. 2001) and excellent thermal conductivity (Xiao et al. 2004), superior to CNTs (Ciofani et al. 2010), a Young’s modulus in the TPa range (Tiano et al. 2014) and they are electrically insulating with a bandgap of ∼5-6 eV (Tiano et al. 2014; Blase et al. 1994). Analogously to CNTs, BNNTs are composed of hexagonal B-N bonds with a partial ionic character and almost the same atomic spacing (Ciofani et al. 2009b; Tiano et al. 2014). The nature of the piezoelectricity of single-walled BNNTs has been demonstrated experimentally, although it is not completely understood (Tiano et al. 2014). Numerical and molecular mechanics simulations, however, have been able to approximate the resultant dipolar vector in single-walled nanotubes after an applied stress (Tolladay et al. 2017; Tiano et al. 2014), while others have found theoretical piezoelectric response values higher than those of piezoelectric polymers (Dai et al. 2009). Likewise, multi-walled BNNTs have also shown experimental signs of piezoelectricity (Bai et al. 2007). Although these characteristics are especially attractive for the aerospace or energy generation industries (Tiano et al. 2014), their applications in the biomedical field have been generally unexplored (Ciofani et al. 2010; 2009b).

Commercially available BNNTs are synthesized by a high temperature-pressure method (Tiano et al. 2014) and come in the form of a puffball of very low density. These nanotubes were characterized by transmission electron microscope (TEM), as shown in **Figure 2A**. This characterization revealed an inhomogeneous mixture of BNNTs with some impurities in the form of hexagonal boron-nitride (h-BN) nanoparticles, as reported by the manufacturer (Tiano et al. 2014). The nanotubes were in general straight and organized in bundles, with diameters as small as tens of nanometers (inset). While one of the key aspects of BNNTs is their chemical stability, this can make their dispersion in aqueous solutions difficult to achieve, hindering their applications in biomedicine (Ciofani et al. 2009b). To improve the dispersion, previous reports have used non-covalent wrapping of BNNTs in poly-L-lysine (Ciofani et al. 2009a; 2010) or glycol chitosan (Ricotti et al. 2013). Here, we chose an alternative method based on dispersion in organic solvents, followed by subsequent solvent transfers until achieving a stable dispersion in ethanol. Firstly, we dissolved the foam-like material in dimethylformamide (DMF) at 0.5 mg/mL under vigorous stirring overnight, until obtaining a brown solution with small aggregates. Then, the samples were sonicated for at least 6 h in an ice-cold bath, until all the aggregates were completely dissolved (Ricotti et al. 2013). Afterwards, the sample was centrifuged and resuspended in a solution of 1:1 of ethanol and DMF at the same concentration and sonicated in cold until achieving dissolution. Finally, the sample was again centrifuged and resuspended in pure ethanol at 0.5 mg/mL. The dispersion was found to be stable for several weeks in ethanol at 4 °C, although small aggregates were formed. Before mixing with the cell-laden hydrogel, the ethanol dispersion of BNNTs had to be sonicated in a cold bath for until the dispersion was homogeneous again, from 10 min to 2 h, depending on the time that the sample had been stored until used.

**Figure 2.**
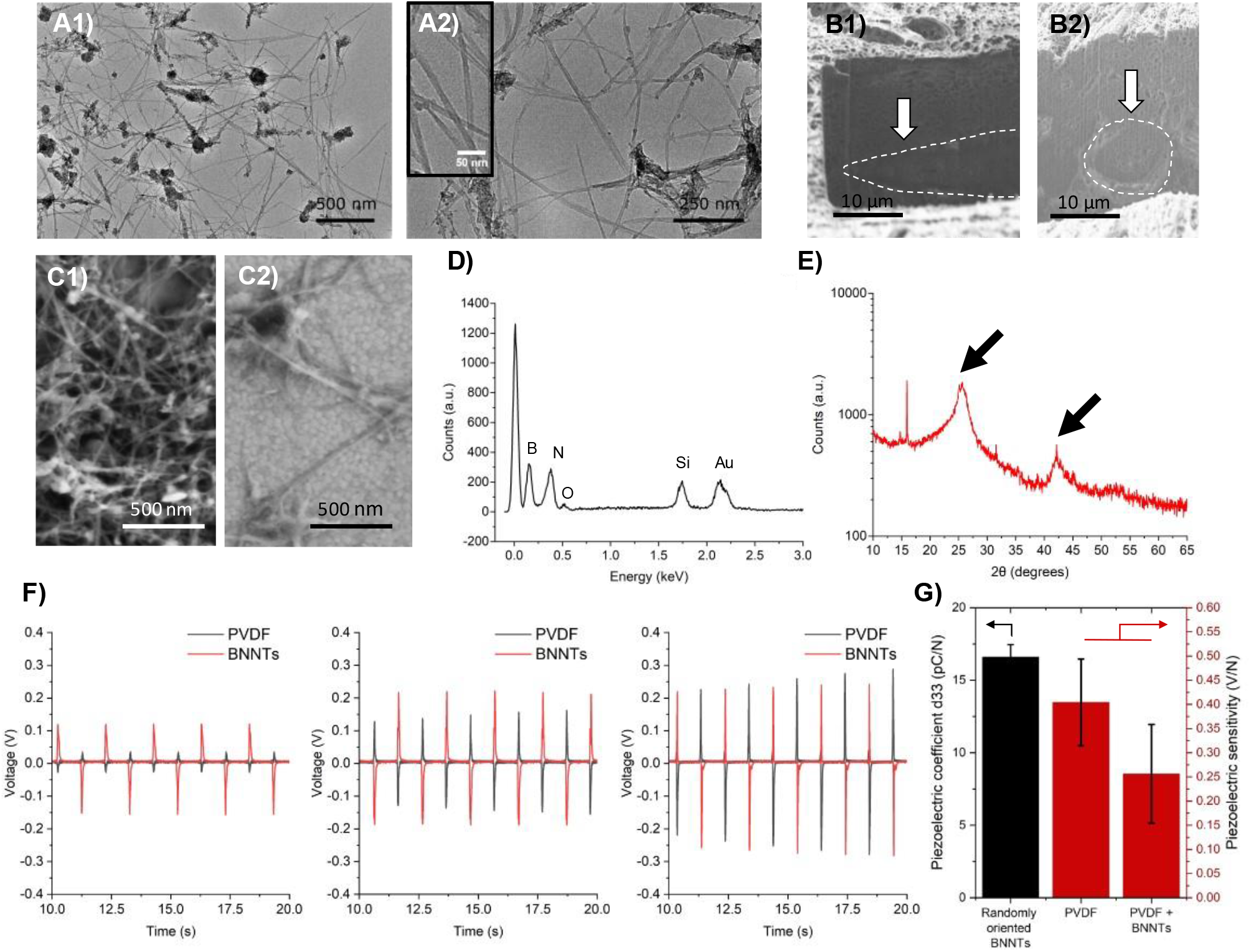
Characterization of BNNTs. A) TEM images at different magnifications of a dispersion of BNNTs. Darker aggregates correspond to boron nanoparticle impurities. B) SEM imaging of a cell-laden hydrogel after performing a cut with FIB in a longitudinal (B1) and transversal (B2) direction, with dashed lines delimiting the shape of myotubes. C) SEM imaging of a sample of BNNTs; and D) EDX performed in the sample of the SEM image where the boron (B) and the nitrogen (N) from the BNNTs can be seen as well as the silicon (Si) and gold (Au) from the base chip. E) XRD pattern of a BNNT sample showing the two peaks, representative of this type of sample. F) Piezoelectric measurements of PVDF membranes with and without BNNTs at three different forces (0.15 N, 0.3 N, 0.75 N), with generation of voltage due to piezoelectricity. G) Effective piezoelectric coefficient (d33) and piezoelectric sensitivity of different types of samples measured with a piezometer and dynamometer, respectively. For randomly oriented BNNTs, the error bars indicate the measurement uncertainty, and for the remaining bars the fitting error.

The disposition on the BNNTs within the matrix and the orientation with respect to the cells is an important parameter to consider: an oriented conformation should induce a higher electrical stimulus and therefore a bigger stretch on the cell. For this, scanning electron microscopy (SEM) images and energy-dispersive X-ray (EDX) characterization of the bio-bots and BNNTs in different matrices were performed. A cut with focused ion beam (FIB) was first delimited on some bio-bot samples (**Figure 2B** and **Figure S1**). The maximum depth achieved with this cut was of 20 µm. Using SEM, the cells disposed in the outer part of the hydrogel and their orientation could be seen, although the inner part of the bio-bot could not be reached. In the longitudinal cuts, cells could be perceived alongside the porous hydrogel (**Figure 2B1**) whereas in the transversal cuts, it was the frontal, yet rounder, part of the muscle fiber that was seen (**Figure 2B2**), indicating the orientation of the muscle fibers (see Supplementary Information for more images). However, due to the difference in between the BNNTs’ diameter and the cells’ (micrometers to nanometers size) the former could not be distinguished from the matrix and the surroundings of the cells neither through SEM imaging nor by performing EDX measurements.

SEM and EDX measurements were performed on samples without cells: first, on collagen hydrogel with different concentrations of BNNTs (5, 10, 15, 20 µg/mL), then inside electrospinned PVDF membranes with 5 µg/mL BNNTs. No BNNTs could be observed in any of the samples, probably due to the dispersion within the matrices and the tiny size of BNNTs. To corroborate with TEM and to further confirm the composition of the nanocomposites, SEM/EDX imaging of a pristine BNNT sample was performed (**Figures 2C** and **D**). Boron (B) and nitrogen (N) peaks could be distinguished perfectly from the sample. Silicon (Si) and gold (Au) peaks, coming from the Au-coated silicon chip used as substrate to deposit the BNNT sample, can also be seen in the spectrum. Finally, **Figure 2E** shows X-ray diffraction (XRD) patterns of pristine BNNTs showing peaks at 25 and 42 degrees, in agreement with the literature (Ferreira et al. 2011).

In order to characterize the piezoelectric properties of the BNNTs and the composite matrix of the bio-bots, two different measurements were performed: dynamometer and piezometer measurements. In the former, the sample is pressured between two clamps that serve as electrodes. There, where the force is applied, the electrodes measure the voltage produced in between the electrodes and translate it to a piezoelectric coefficient. The piezoelectric coefficient of pristine BNNTs was measured with the sample as purchased, obtaining a value of d33 = 16.6 ± 0.9 pC/N for randomly oriented BNNTs and confirming the piezoelectric behavior of the nanocomposites, as per the literature on the topic (Tiano et al. 2014). Hydrogel samples with BNNTs at 5 µg/mL concentration, with and without cells, were measured but the value obtained was not stable, mainly due to the fragility, heterogeneity, and thin thickness of the hydrogel once dehydrated. In order to completely prove their piezoelectric effect, we also measured the piezoelectric coefficient of BNNTs in a composite matrix of PVDF nanofibers.

To obtain a more precise measurement with aligned BNNTs, dynamometer measurements were performed on samples made of electrospun PVDF nanofibers (which are known to be piezoelectric), with and without BNNTs (see Methods section for more details). As reported seen in **Figure 2F**, both samples display piezoelectric behavior, as the voltage generated increases when the value of the applied force increases. At the smallest forces (0.15 N), the samples with BNNTs had a better electrical performance than the control ones; but, as the force increased, the signal produced by the PVDF membranes without BNNTs equated the ones with BNNTs at larger forces (0.75 N) and thus their sensitivity decreased. This effect suggests two hypotheses: i) the piezoelectric coefficient of BNNTs is of the opposite sign that PVDF and decreases its piezoelectric sensitivity; and ii) the electric outputs can be enhanced due to the addition of the BNNTs not only due to the piezoelectric effect, but to the expected increase in the dielectric constant and Young’s modulus of the nanocomposite (H. S. Kim et al. 2018). The voltage jump between the different sets of forces is reduced for the BNNT+PVDF sample, indicating that the nanocomposites have reduced the magnitude of the piezoelectric coefficient of the sample. This matches previous reports in the literature, in which the piezoelectric coefficient of PVDF has been reported to be negative (H. S. Kim et al. 2018; Katsouras et al. 2016; Dahiya and Valle 2013; Bowen et al. 2014; Nix and Ward 1986) and that of BNNTs to be positive (Tiano et al. 2014). In this scenario, this leads to the equilibrium of the dipoles until one reaches the potential produced by the other. We performed FEA simulations (**Figure S2**) to prove this hypothesis. We simulated a PVDF fiber with a BNNT inside of it and we applied a compressive force at each extreme of the fiber. The interplay between piezoelectric effects of different sign indeed shows how the overall piezoelectric coefficient of the PVDF+BNNT fiber is reduced in magnitude, supporting the hypothesis of the reduced effect observed in the sensibility experiments (**Figure 2G)**.

### Physical and biological characterization of BNNT-loaded bio-bots

Force measurements on control and BNNT-loaded skeletal muscle tissues were performed by deflection of two PDMS posts as previously reported (Mestre, Patiño, Barceló, et al. 2019). Tissue constructs were fabricated following protocol outlined above and then transferred into the two-post system for differentiation (**Figure S3**). After 3 and 5 days in DM, absorbance values after performing a PrestoBlue viability assay were within normal range and without differences between control and BNNT samples, indicating no decrease of viability due to the addition of the nanocomposites (**Figure 3A**). Immunostaining of MyHCII and cell nuclei showed no differences between BNNT and control samples, since both of them exhibit elongated and well-aligned myotubes (**Figure 3C**), as previous reports in the literature also showed (Ciofani et al. 2010). The effects on force generation and bio-bot performance, hence, could come from differences at the protein level, such as remodeling of the sarcomeric structure.

**Figure 3.**
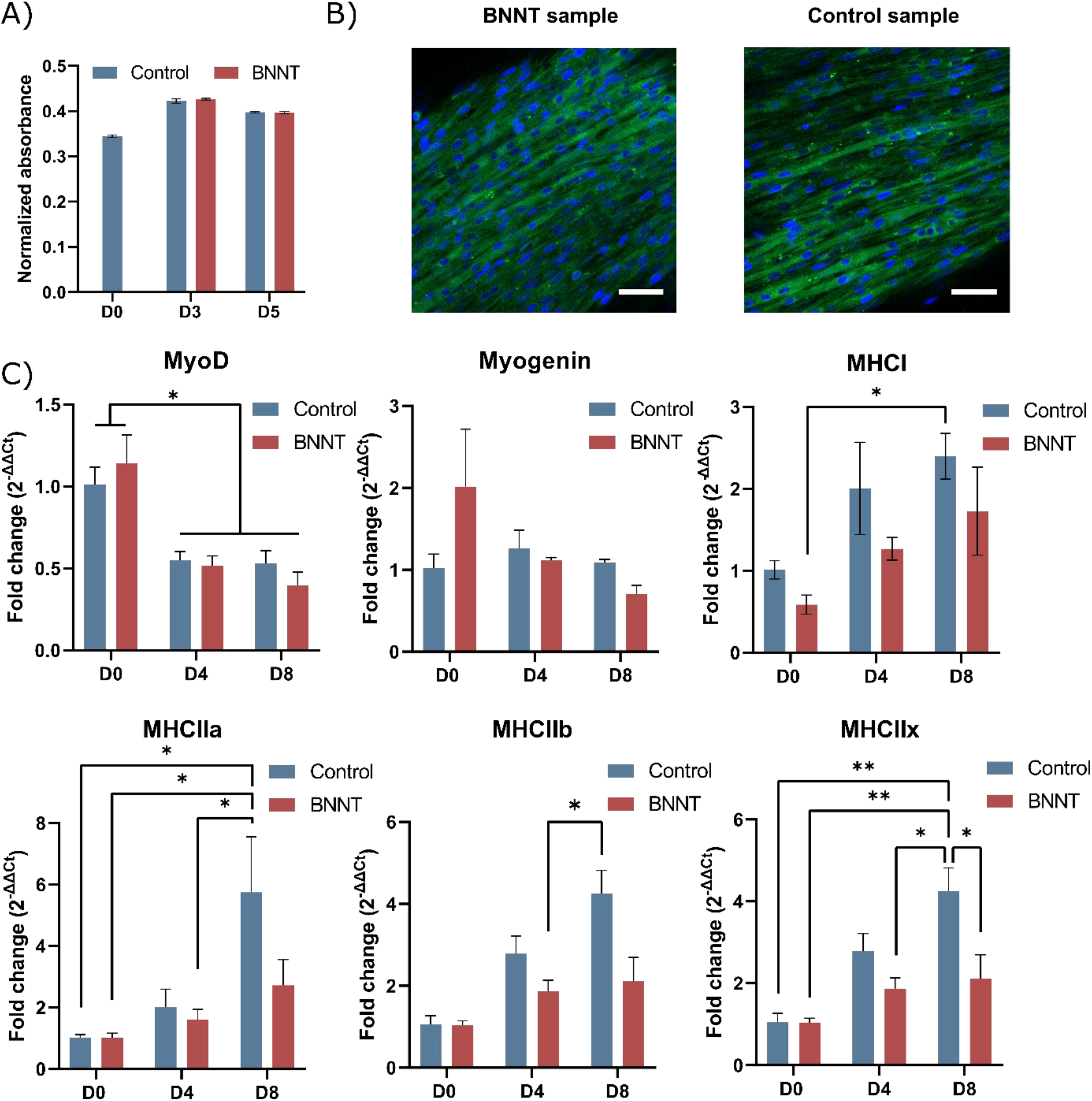
Biological characterization of control and BNNT-loaded bio-bots. A) Normalized absorbance of PrestoBlue viability assays of the two types of bio-bots, showing no difference in viability after the addition of nanocomposites. B) Immunostaining of MHC of BNNT and control samples, showing well differentiated and aligned myotubes in the hydrogel. C) Results of qRT-PCR for six genes of interest, relating to differentiation (MyoD and Myogenin) and maturation (MHCI, MHCIIa, MHCIIb and MHCIIx) of skeletal muscle tissue. Error bars indicate standard error of the mean. * p < 0.05. Two-way analysis of variance (ANOVA) was performed followed by Tukey’s post hoc test for ultiple co pariso s. Shapiro-Wilk test was performed to assess normality of the model residuals.

To assess the effect of BNNTs on mRNA maturation, real time quantitative polymerase chain reaction (RT-qPCR) was performed at different time points (day 0, 4 and 8 of differentiation) to evaluate the expression of differentiation markers, such MyoD and Myogenin, as well as the maturation-related genes MyHCI, MyHCIIa, MyHCIIb, and MyHCIIx (**Figure 3C**). We observed a down-regulation of MyoD and Myogenin expression overtime, especially in BNNT-loaded bio-bots, as expected during the first days of differentiation. The four isoforms of MyHC were more expressed at the late stages of differentiation, when the muscle starts maturation and multinucleated myotubes with internal sarcomeric structure and contractility are formed (Cusella-De Angelis et al. 1992). However, the up-regulation of the maturation-related genes was not statistically significant in BNNT-loaded bio-bots, suggesting that the terminal state of maturation was achieved, leading to a halt in protein synthesis along with a stabilisation of MyHC mRNA and protein levels (DeNardi et al. 1993). Such event was also observed in a previous report where skeletal muscle-based actuators undergoing long term electrical stimulation showed improved force output while down-regulation of maturation-related genes (Mestre, Patiño, Barceló, et al. 2019). Finally, in comparison with control samples, no significant differences in gene expression at the same differentiation day where observed, indicating that BNNTs did not show negative effects on gene expression during muscle differentiation and maturation.

### Motion performance of BNNT-loaded bio-bots

Finally, the performance of BNNT-loaded bio-bots in terms of speed was compared to control bio-bots at day 6 of differentiation. **Figure 4A** shows two illustrative examples of a bio-bots walking upon electric pulse stimulation (EPS). The tracking algorithm described in the methods section was applied to follow a corner in the skeleton of the bio-bots and their speeds were computed from the trajectory. Despite showing great variability between samples, BNNT-loaded bio-robots showed generally higher speeds than control bio-bots, with values comparable to those reported previously (Cvetkovic et al. 2014). In particular, the average speed at 1 Hz was slightly higher for BNNT bio-bots but the speed at 4 Hz was greatly increased, showing statistically significant differences compared to the control cases (**Figure 4B**). The best two cases of motion for each kind of bio-robot are shown in **Figure 4C**, where a fitting was performed to obtain their speeds. It can be seen how BNNT-loaded bio-bots could reach speeds up to 234 μm/s, three times higher than the control cases at roughly ∼80 μm/s. Videos of the best performing BNNT and control bio-bots at 2 and 4 Hz can be seen in **Videos S1-4** of the Supplementary Information. **Figure 4D** displays the tracked trajectories of the examples of **Figure 4C**, where it can be observed how they are completely directional. However, several of the bio-bots showed significant rotational motion, probably due to inhomogeneities in the skeleton during 3D printing and in the muscle itself. These bio-bots, therefore, were not included in the speed calculations, but only directional ones. We believe this increase in speed to be due to a greater force production of the bio-bots. At day 6 of differentiation, control samples showed forces in the range of ∼80 μN (**Figure 4E**), similar to previous reports (Cvetkovic et al. 2014; Raman et al. 2016). However, BNNT-loaded muscle tissues showed more than a two-fold increase in their force generation, reaching values of 250 μN, with statistically significant differences.

**Figure 4.**
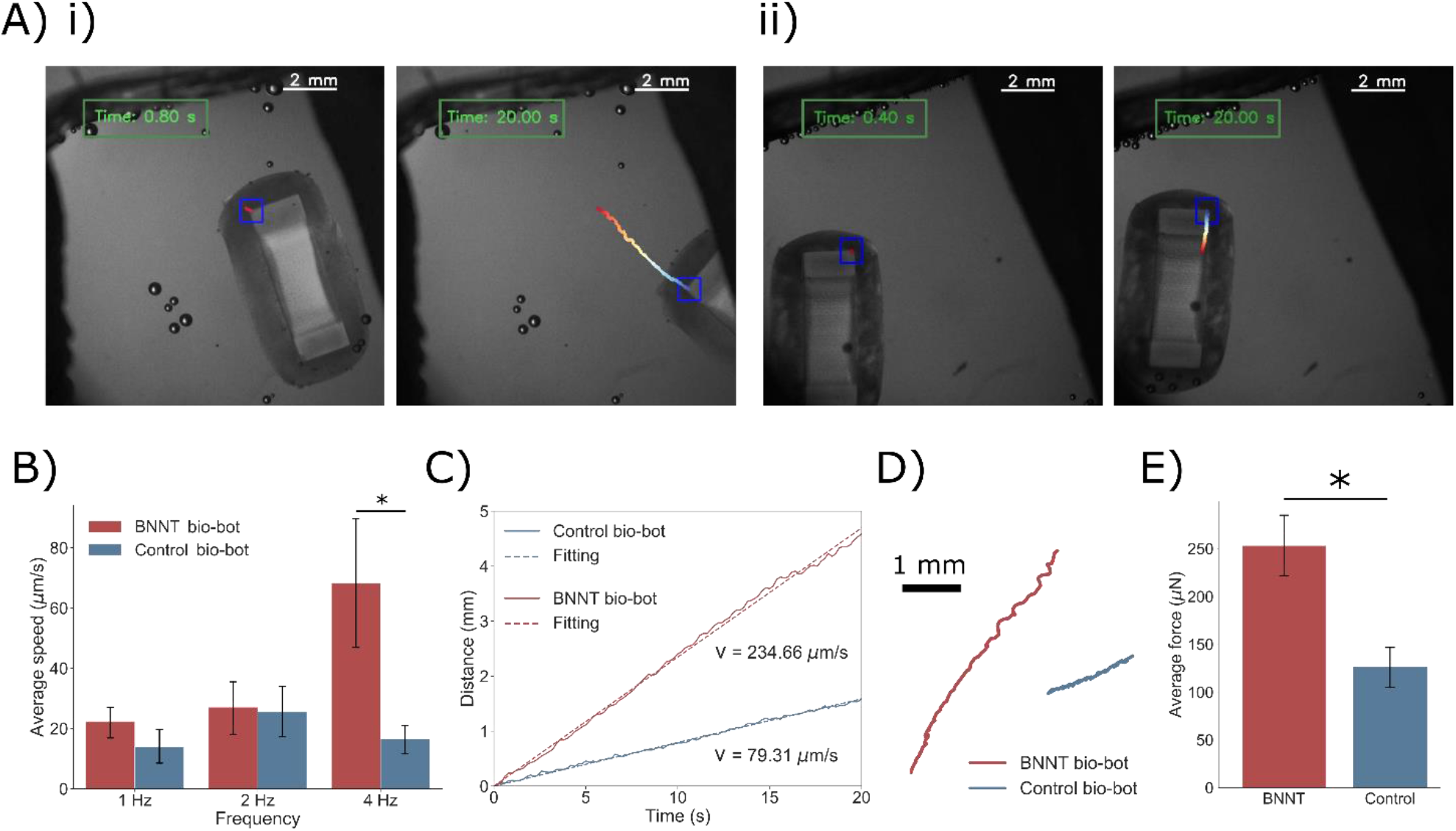
Motion analysis and comparison of control and BNNT bio-bots. A) Snapshots of a tracking of a (i) BNNT-loaded bio-bot and a (ii) control bio-bot moving upon EPS. B) Average speeds reached by control and BNNT bio-bots according to different frequencies. C) Examples of the two best cases of motion of control and BNNT bio-bots for a period of 20 s at 4 Hz, and D) their trajectories. E) Force measurement of BNNT and control samples at day 6 of differentiation (t-test with * p < 0.05).

To understand the synergy of multiple effects produced at the micro- and nanoscale and how it translates into the macroscopic effect of the displacement of the bio-bot, we computed a simulation of a hydrogel-muscle matrix with several BNNTs distributed inside of it (**Figure 5** and **Figure S4**). Compression pressures in the range of 0.001 to 10 kPa were simulated at both ends of the hydrogel-muscle structure. This range of values matches the typical tensions exerted by the passive compaction of the tissue during differentiation and the active contraction of the myotubes, as reported in the literature (Sakar et al. 2012; Raman et al. 2016). The simulation results show that, as expected, the piezoelectric BNNTs embedded in the matrix translate the stress of the compression into electric field to the surrounding myotubes. As can be seen in **Figure 5A**, the surface stress is especially high at BNNT surface because of their small size, and thus the electric field due to piezoelectricity (**Figure 5B**) is also greater near the surface of the nanocomposites. Most importantly, the combined effect of all the BNNTs makes the electric field within the hydrogel be non-zero, indicating that the myotubes are exposed to a positive electric field magnitude (**Figure 5C**), which helps in their differentiation process, supporting our initial hypothesis. Finally, **Figure 5D** show the electric field on a line longitudinally along the hydrogel-muscle matrix for pressures in the range of 0.001-10 kPa. As expected, the larger the compression force, the larger the electric field magnitude. Although the electric field magnitude some radii away from the BNNTs might seem small compared to their interface, this value is still high enough to produce a significant response in the myotubes within the matrix, since during external EPS typical electric fields are in the range of 1 V/mm or lower (Ito et al. 2014).

**Figure 5.**
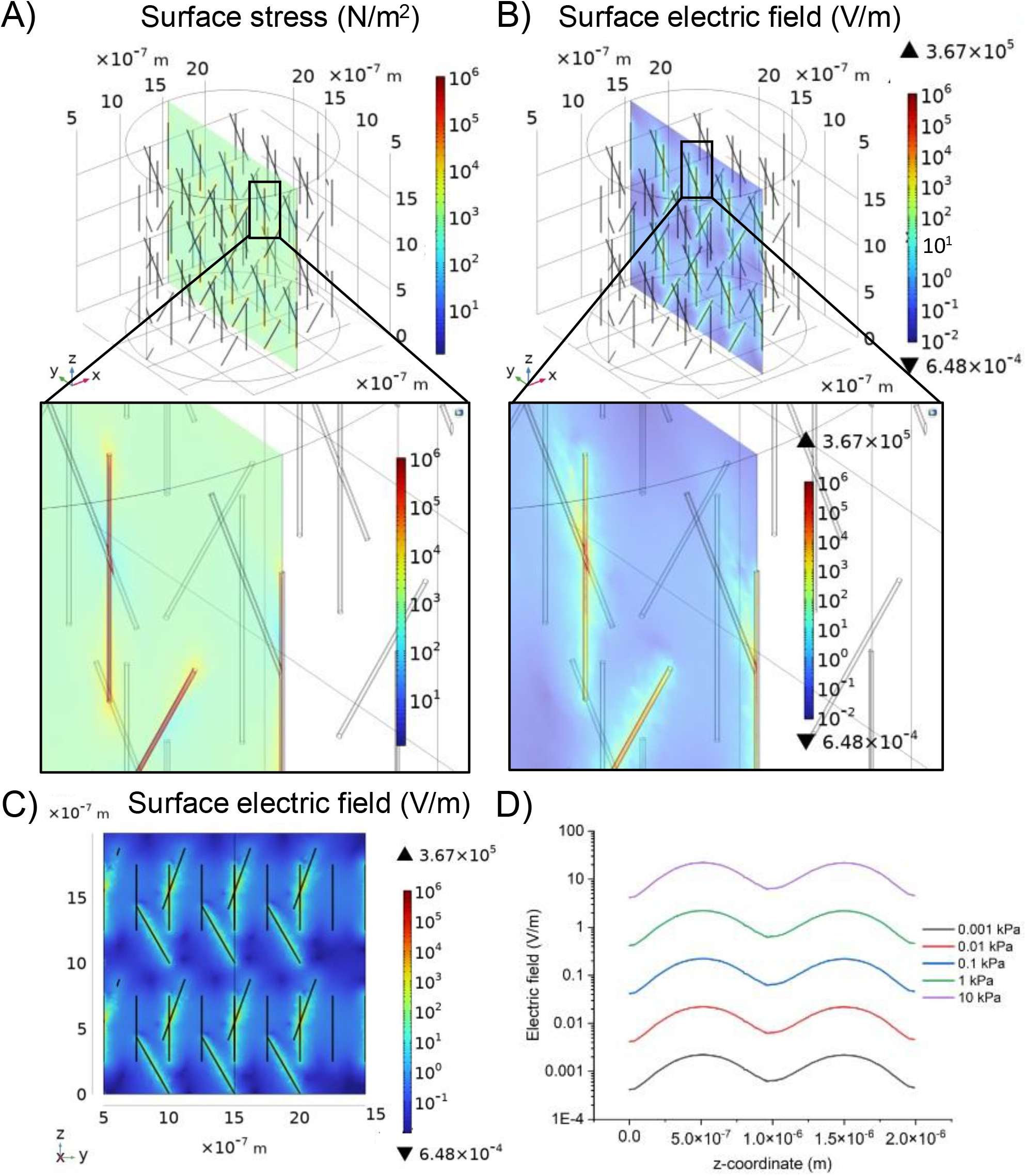
COMSOL Multiphysics simulations of a matrix of hydrogel with BNNTs embedded in it. A) Surface stress across a cut in the matrix after the application of a compression force of 1 kPa, and B) the surface electric field generated by the piezoelectric coefficient of the BNNTs, and C) a surface plot of the same case. D) Distribution of the electric field along a longitudinal line (z-axis) for different compressive tensions. The oscillations come from neighboring nanocomposites, where the electric field is stronger.

## Conclusions

We have demonstrated the integrating piezoelectric BNNTs into the cell-laden hydrogel of walking bio-bots. A dispersion of BNNTs was obtained by dissolution in DMF and several solvent changes until reaching a stable solution in ethanol. Bio-bots loaded with BNNTs showed enhanced motion, as demonstrated by tracking of bio-bots actuated by EPS. Moreover, we demonstrated that these improvements in the motion were a result of a stronger force output of the nanocomposite-laden muscle tissue. By performing force measurements, we found that the average force of BNNT-loaded tissues showed a 2.5-fold increase with respect to control samples. Results from the genetic characterization of the bio-bots showed that the expression of the maturation markers was lower in the BNNT-loaded samples, although not statistically significant in general. However, the enhanced force contraction produced by these BNNT-loaded tissues indicates that the muscle fibers are fully matured and contractile, suggesting they have reached terminal maturation. Characterizations and FEA simulations demonstrate that the BNNTs show piezoelectricity and that a random distribution of BNNTs inside the hydrogel can produce a non-zero electric field in the muscle cell membrane. This supports our hypothesis that integration of nanocomposite into muscle-based bio-bots can improve their performance in terms of motion and force output.

These preliminary results pave the way was towards a more complex actuation of bio-hybrid robots by the integration of nanocomposites with different capabilities, in this case, piezoelectric nanoparticles that improve the performance of the tissue. Future studies should investigate the piezoelectric effect for remote actuation of these bio-bots by the application of ultrasonic waves. Other nanocomposites could be also combined to boost the performance and efficiency, such as magnetic nanoparticles in the cell-laden hydrogel or the artificial scaffold, to guide the bio-hybrid robots through magnetic fields or graphene-based nanocomposites that could improve the mechanical and conductive properties of the hydrogels. The multidisciplinary integration of recent advances in smart materials, nanotechnology, tissue engineering and soft robotics will be key for the future development of advanced bio-hybrid robotics

## Materials and methods

### Bio-bot skeleton fabrication

PEGDA with a number average molecular weight (Mn) of 700 (Sigma-Aldrich) was mixed with PEGDA 575 Mn in a 1:1 ratio. A solution of PEGDA 700/575 at 16% v/v in ultrapure water with lithium phenyl-2,4,6-trimethylbenzoylphosphinate (LAP; Sigma-Aldrich) at 0.1% wt/v was prepared. A red dye, Sudan I (Sigma-Aldrich) was included in the solution at a 0.15% wt/v and sonicated until fully dissolved. The skeletons, based on previous CAD designs (Cvetkovic et al. 2014), were 3D printed with the PICO2 SLA 3D printer from Asiga. Sudan I dye was necessary to improve the definition of the structures by reducing the light scattering effects. After printing, the skeletons were soaked in 10% bleach to remove the dye, left in isopropanol overnight and then maintained in PBS until use.

### Cell culture

C2C12 mouse myoblasts were purchased from ATCC. Growth medium (GM) consisted of high glucose Dullbecco’s Modified Eagle’s Medium (DMEM; Gibco®) supplemented with 10% Fetal Bovine Serum (FBS; ThermoFisher), 200 nM L-Glutamine and 1% Penicillin/Streptomycin (Lonza). Cells below passage 6 were used before reaching 80% confluency in T-75 flasks. Differentiation medium (DM) consisted of high glucose DMEM containing 10% Horse Serum (Gibco®), 200 nM L-Glutamine (Gibco®), 1% Penicillin-Streptomycin (Gibco®), 50 ng/ml insulin-like growth factor 1 (IGF-1, Sigma-Aldrich) and 1 mg/ml 6-aminocaproic acid (ACA, Sigma-Aldrich).

### Fabrication of control and BNNT-loaded bio-bots

Myoblast-laden hydrogel were composed of Matrigel® (ThermoFischer), fibrinogen (Sigma-Aldrich) and thrombin (Sigma-Aldrich) in GM supplemented with ACA. Cells at 80% confluence were harvested by adding 3 mL of TrypLE Express (ThermoFischer) for 5 mL and then neutralized with 3 mL of GM. Then, cells were centrifuged at 300g for 5 min and separated into 15-mL Falcon tubes with 3 million cells in each on. For cell encapsulation, 115 μL of GM + ACA were added to a pellet with 3 million cells and homogenized thoroughly. Then, 6 μL of a 100 U/mL solution of thrombin and 90 μL of Matrigel® were added and homogenized. Finally, 75 μL of stock solution of fibrinogen at 16 mg/mL was added, homogenized fast, and 120 μL of solution was casted in one injection mold, preparing a total of two each time. The tissue constructs were left in a cell incubator at 37 °C and 5% CO2 atmosphere for 1 h until Matrigel® was crosslinked and GM supplemented with ACA was added.

BNNTs (product code R2-beta) were purchased from BNNT, LLC. The material came in the form of a puffball and small quantities of it were dispersed in DMF at a concentration of 0.5 mg/mL, as described in the main text, by first leaving it under vigorous stirring overnight and then sonicating in ice for at least 2 h. The sample was centrifuged at 1400 rpm in Eppendorfs and the solvent was changed to DMF and ethanol in a 1:1 ratio, maintaining the same concentration of BNNTs. The process of sonication and centrifuging was repeated until obtaining a solution in pure ethanol. After solvent change to ethanol, the sample could be stored at 4 °C in cold. Before mixing with the hydrogel, the dispersion was again sonicated in cold for at least 1 h or until it was completely homogeneous and without aggregates. Considering that the total amount of hydrogel needed was 286 μL (as per last paragraph’s quantities), 2.86 μL of the stock solution of 0.5 mg/mL BNNTs in ethanol was added to achieve a final concentration of 5 μg/mL in the hydrogel. The addition of the BNNT dispersion was performed right before addition of fibrinogen and homogenized thoroughly, without observing presence of aggregates.

To stain BNNTs with Rhodamine, a 1mg/mL BNNTs solution was mixed with 10ug/mL of Rhodamine at a ratio of 1:4. After overnight incubation at room temperature with agitation (300 rpm), stained BNNTs were washed 3 times via suspension in ethanol and centrifugation for 5 min at 13000 rpm.

### Biobot tracking

The script for tracking the motion of the bio-robots was developed in Python 3.7 using the library OpenCV (v. 4.1.2). Videos recorded with a microscope camera were loaded into the script using the VideoCapture function. The first frame of the video was prompted, and the user had to select an ROI around the bio-robot with the SelectROI function. A BOOSTING tracker type was created (function TrackerBoosting_create) by initializing it with the selected ROI and it was then applied throughout the whole length of the video, frame by frame. The central position of the ROI was stored, and the trajectory was added in the video with a colour code to indicate time. Finally, a linear fitting of the form *x* = *v·t* was applied to the displacement *vs* time data to obtain the speed of the bio-robot.

### Motion analysis of bio-bots and force measurement of muscle tissue

Motion characterization was performed under an up-right microscope with a set of home-made stimulation electrodes composed of two platinum rods (Figure S5). A waveform generator with an amplifier (AD797) were used to apply squared biphasic pulses by connecting a capacitor in series to minimize electrolysis created by the Pt electrodes. Bio-bots were recorded at different frequencies after placing them in fresh, warm DMEM and the motion was analyzed with the home-made tracking script described in the previous section.

Force measurement was performed as described in (Mestre, Patiño, Barceló, et al. 2019; Mestre, Patiño, Guix, et al. 2019). Briefly, control and BNNT-loaded tissues were transferred into a 2-post system 3D-printed with PDMS instead of into a bio-bot skeleton and they were left to differentiate for 6 days in DM supplemented with ACA. Force measurement was done at 1 Hz and recorded under an inverted microscope, following our previously reported protocol (Mestre, Patiño, Guix, et al. 2019).

### Immunostaining

Tissue constructs were washed three times in PBS and then fixed with a 3.7% paraformaldehyde in PBS for 15 min at RT, followed by three washes in PBS and stored until use. For immunostaining, cells were permeabilized by 0.2% Triton-X-100 in PBS. After washing thrice in PBS, the constructs were incubated with 5% Bovine Serum Albumin (BSA) in PBS (PBS-BSA) to block unspecific bindings. Then, the tissues were incubated overnight at RT and in dark conditions with a 1/400 dilution of Myosin 4 Monoclonal Antibody (MF20; eBioscience) as primary antibody and 4′,6-diamindino-2-phenylindole (DAPI; Sigma-Aldrich) at 1:5000. The next day, the samples were washed thrice in BPS and Alexa Fluor® 568 was added for 2 h at RT in dark as secondary antibody. The tissue samples were observed under CSLM (Zeiss).

### PCR

Total RNA content was extracted from 3 biological replicates using RNeasy® Plus Mini kit (#74134, Qiagen) following manufacture’s protocol. Extracted RNA was quantified by absorbance at 260 nm in a nano-drop spectrophotometer (ND-1000, Thermo Scientific™). The cDNA was synthesized from 500 ng of RNA using the RevertAid First Strand cDNA Synthesis Kit (Thermo Scientific™, K1621). RT-qPCR reactions were performed using PowerUp SYBR Green Master Mix (Applied Biosystems, A25742), following to manufacturer’s instructions, with 500 ng of cDNA and the target primers in a total volume of 10 µl in a StepOnePlus Real-Time PCR System (Applied Biosystems, 4376600). All genes were normalized to the expression levels of GAPDH. Melt-curve analyses were performed to ensure that only one amplicon was being produced.

The following primers were used:

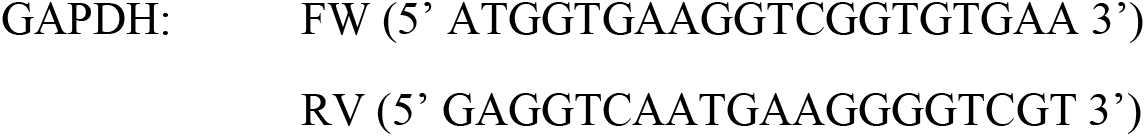
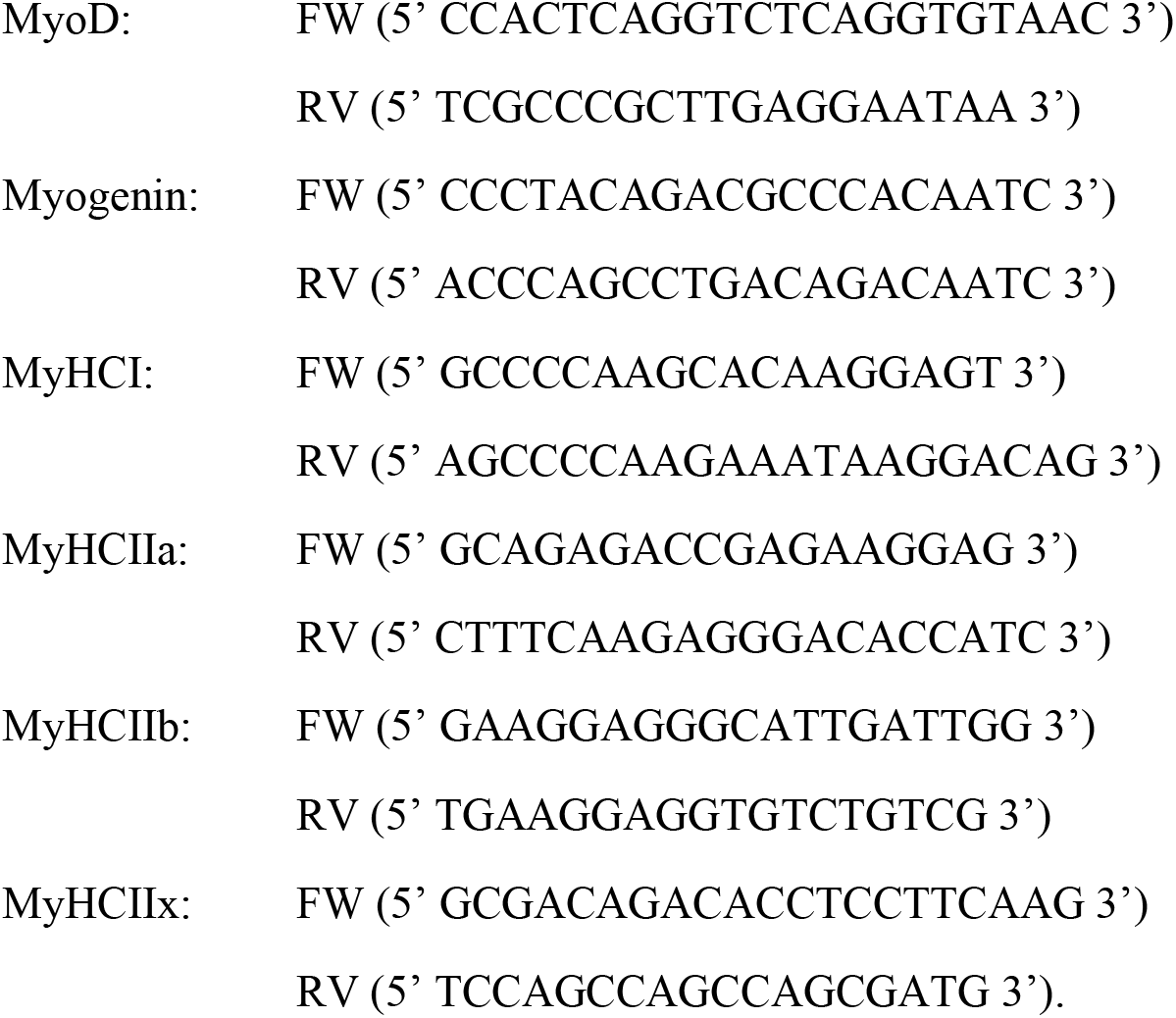

### Viability

Cell viability was evaluated with PrestoBlue™ cell viability reagent (Thermo Scientific™, A13262) following manufacturer’s protocol. PrestoBlue™ reagent was added at a ratio of 1:10 with respect to the total volume of cell medium and samples were incubated for 2 h at 37°C. Then, 50 µL of media were taken and absorbance was measured at 560 nm in a Benchmark Plus Microplate Reader, normalized to the value at 600 nm, subtracting the background of only-medium controls.

### Simulations

Finite element simulations using the integrated electrostatic physics of COMSOL were performed to simulate a muscle matrix –composed by several muscle fibers in a syncytium-embedding BNNTs (Figure S4). BNNTs were defined as piezoelectric with the following parameters: Young modulus 1180 GPa (Trivedi, Sharma, and Harsha 2014), relative permitivitty 3.29 (Laturia, Van de Put, and Vandenberghe 2018) and d33 0.76 C/m^2^ (Sai and Mele 2003; Salehi-Khojin and Jalili 2008). A similar design with these same parameters was used for the simulation in supplementary material (Figure S2) where a single BNNT embedded in a PVDF fiber is compressed. A compressive force in opposed direction in the z-axis was applied in both faces of the cylinder to generate the compressive stress on the muscle syncytium. The study performed was a parametric study applying pressures from 0.001 to 10 kPa. The electric field produced by these forces was studied and compared among each other. Then, a point was selected in the simulation to evaluate the electric field along the muscle in the z-axis to justify that the electric field was non-zero. The mesh used for this simulation was anisotropic as it allowed the accurate representation of smaller elements –BNNTs- in the thinner domains and larger elements –muscle- in the system. A similar simulation was performed mimicking the stretching movement of the muscle by changing the direction of the forces applied on the cylinder bases in the z-axis.

### FIB and SEM/EDX imaging

Using a SEM Auriga-40 (Carl Zeiss), samples were characterized in order to see the relation between the collagen fibers, the BNNTs and the cells within the bio-bots; and C2C12 mouse myoblasts morphology inside the hydrogel. Previously, a vertical and a horizontal section was performed using a FIB (Crossbeam 1560XB, Carl Zeiss) of approximately 20 µm depth. Images were taken with a magnification ranging from 500 to 100K X at 2 keV of energy with the SE2 detector. To maintain the cell and the hydrogel structure, the samples were fixed with paraformaldehyde 4% and dehydrated following an increasing concentration of ethanol (50, 70, 90, 100, 100 %). Finally, the critical point of the sample (BAL-TEC CPD030), and a metallization (BAL-TEC MED 020) with a carbon pellicule were performed.

Other measurements with SEM Auriga-40 (Carl Zeiss) include the imaging and EDX (INCA Energy Operator) of a sample of BNNTs dissolved in pure ethanol. These were deposited on top of a silicon chip with a layer of 50 nm of gold and examined when the ethanol evaporated at RT. Also, images were taken with magnification ranging from 500 to 100K X at 2 and 10 keV of energy with the SE2 detector.

### XRD

BNNTs suspended in pure ethanol were deposited on top of a Si-Au chip, let dry at RT and analyzed using Brucker D8 Advance X-Ray Diffraction (XRD) equipment with a Lynxeye-XE-T detector (Brucker). A Cu radiation source was used, and the X-ray source operated at 40 kV and 40 mA. The Göbel mirror was situated 0.6 mm from the divergence slit, 6 mm from the sample, that was 8 mm from the receiving slit. The spot of the sample was selected using a laser.

### Piezoelectric measurements with dynamometer

A dynamometer conformed by a Mark-10 M5i force indicator and MR03-20 force sensor (Aname) associated to a motorized stage and a stepper motor (LSQ075B-T3-MC03 and X-MCB1-KX13B respectively, Zaber), was used to apply different forces in a range of 0.5 to 3 mm in the z-axis from the sample. The weight moved with a velocity of 20 mm/s. Each sample had a top and a bottom copper electrode connected to the acquisition system (BNC-2110, National Instruments). Voltage data was acquired at a 10000 sampling rate. A filter of 100 Hz was applied using a low-noise preamplifier (Model SR560, Standford Research Systems) with 12 dB low-pass and DC coupling. The whole system was locked inside a Faraday cage to reduce external electromagnetic interferences.

For the preparation of the PVDF matrix, 180 kDa PVDF (Sigma Aldrich) was dissolved in a 20% w/v concentration in a mixed solvent system of DMF/Acetone (4:6 ratio). DMF was purchased from Sigma Aldrich while acetone was provided by BASF. This solution was stirring overnight with a magnet in a hotplate at 60 °C and 740 rpm.

When the PVDF pellet was dissolved, the solution was separated in half and BNNTs suspended in acetone were added in a concentration of 5 µg/mL. The fixed parameters for the electrospinning were as follows: flow rate of 300 µL/h, applied voltage of 18.5 kV, nozzle tip-collector distance of 15 cm, nozzle tip with interior diameter of 0.9 mm, and syringe with interior diameter of 9 mm. PVDF was electrospun on a PET-Au substrate for 20 min with a Fluidnatek LE-10 (Bioinicia) equipment. The morphology and diameter of the fibers were checked using the SEM microscope (Auriga-40, Carl Zeiss). In electrospinning, the diameter of the fiber depends directly on the voltage applied inside the equipment. We expect that the 18.5 kV applied was enough to align the BNNTs inside the fiber. Later, a top copper electrode was added and alligator clips from the dynamometer set-up were connected to the top and the bottom electrode of the membrane. A range of distances were set to apply different forces on the sample, which acted as a beam; and in response to the pressure executed on its tip, the PVDF membrane stretched and compressed their fibers so the BNNTs produced an electrical response to the mechanical stimuli.

### Piezoelectric measurements with piezometer

Using a piezometer YE2730A d33 METER (APC International, Ltd.), a force of 252·10^−3^ N directly to the BNNTs, obtaining the value for d33. The materials were compressed between two steel plates of 100 µm serving as top and bottom electrodes.

### Statistical analysis

For motion characterization, pairs of speeds at the same frequency (control vs BNNT) subtracted from fitting to motion curves were performed a F-test for two-sample variance. As some of the pairs showed unequal variance, a t-test assuming unequal variances was performed (N = 7-14). Significance was set at p-value < 0.05. For force characterization, sets of forces of control and BNNT-loaded samples were performed an F-test for two-sample variance. As the variances were not significantly different, a t-test assuming equal variances was performed (N = 6). Significance was set at p-value < 0.05. For genetic characterization RT-qPCR was performed with N=3 independent repeats. The comparative qRT-PCR statistical analyses were performed using the 2−ΔΔCT method, where all genes were normalized to the expression levels of GAPDH and to the expression levels of control samples at D0. Normality of the model residuals was assessed with Shapiro-Wilk test and difference in gene expression was analyzed with two-way analysis of variance (ANOVA) followed by Tukey’s post hoc test for multiple comparisons. Statistical significance is indicated by p < 0.05.

## Acknowledgements

R. M. thanks “la Caixa” Foundation through IBEC International PhD Programme la Caixa Severo Ochoa fellowships (code LCF/BQ/SO16/52270018 and UK Research and Innovation (UKRI) grant reference MR/S032711/1. M.G. thanks MINECO for the Juan de la Cierva fellowship (IJCI2016-30451), the Beatriu de Pinós Programme (2018-BP-00305), and the Ministry of Business and Knowledge of the Government of Catalonia. The research leading to these results has received funding from the grant RTI2018-098164-B-I00 funded by MICIN/AEI/10.13039/5011000110333 and by “FEDER Una manera de hacer Europa” (BOTSinFluids project), the CERCA program by the Generalitat de Catalunya, the Secretaria d’Universitats i Recerca del Departament d’Empresa i Coneixement de la Generalitat de Catalunya through the project 2017 SGR 1148, and the “Centro de Excelencia Severo Ochoa”, funded by Agencia Estatal de Investigación (CEX2018-000789-S). This project has received also funding from “la Caixa” Foundation under the grant agreement LCF/PR/HR21/52410022 (BLADDEBOTS project). This study was supported by the Spanish Government (PID2020-119350RA-I00 and EUR2020-112082) and La Caixa Foundation under the Junior Leader Retaining program (LCF/BQ/PR19/11700010). The work done at UIUC was funded by NSF EFRI C3 SoRo No. 1830881 and with support from the NSF Science and Technology Center Emergent Behavior of Integrated Cellular Systems (Grant CBET0939511).

