## Supplementary Information for "Improved performance of biohybrid muscle-based bio-bots doped with piezoelectric boron nitride nanotubes"

### Supporting Information

#### Videos

**Video S1 - BNNT-loaded biobot moving at 4 Hz.** Bright-field video of a biobot loaded with BNNT moving upon electrical stimulation of 4 Hz with tracking.

**Video S2 - BNNT-loaded biobot moving at 2 Hz.** Bright-field video of a biobot loaded with BNNT moving upon electrical stimulation of 2 Hz with tracking.

**Video S3 - Control biobot moving at 4 Hz.** Bright-field video of a control biobot moving upon electrical stimulation of 4 Hz with tracking.

**Video S4 - Control biobot moving at 2 Hz.** Bright-field video of a control biobot moving upon electrical stimulation of 2 Hz with tracking.

### Figures

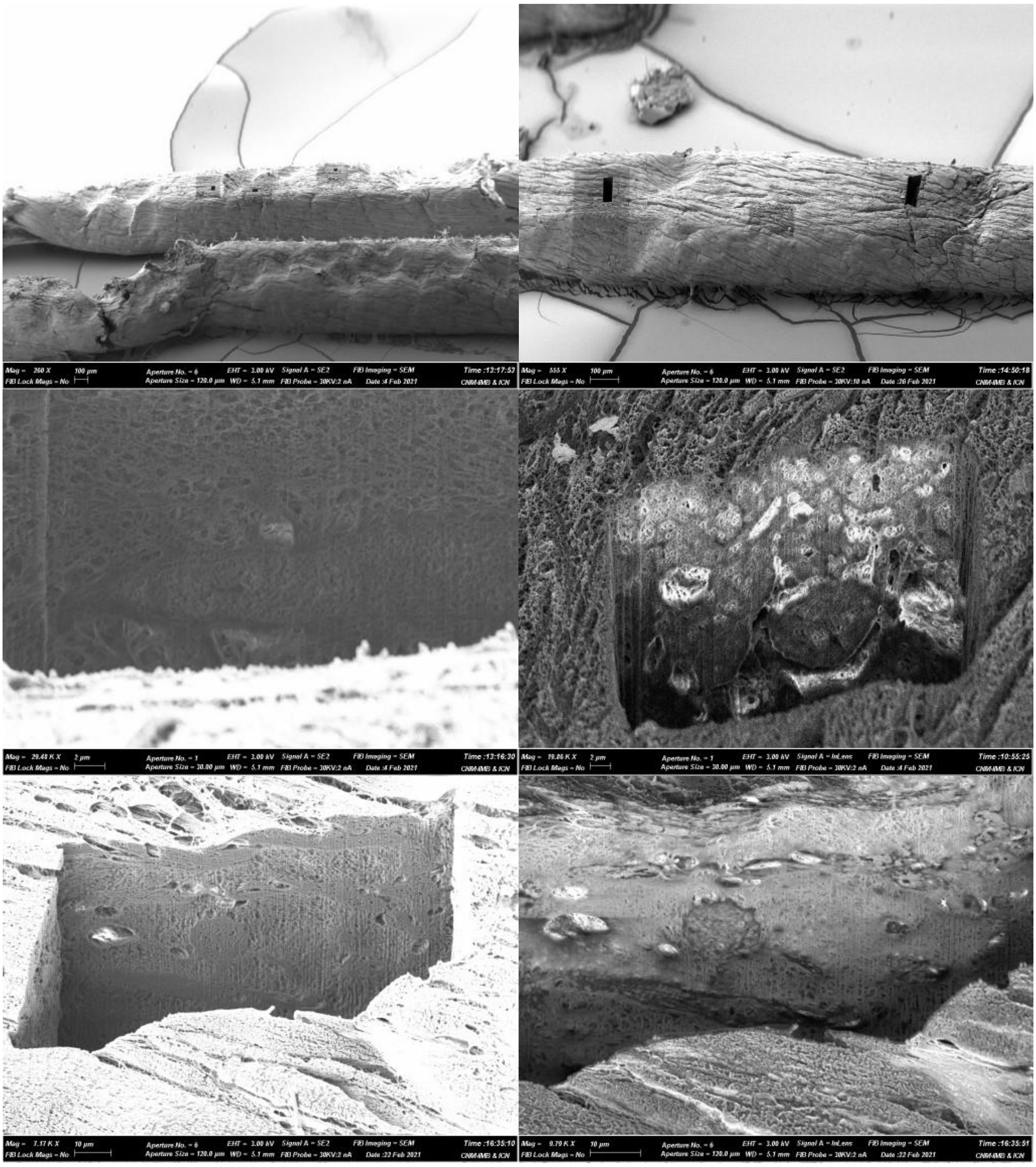

**Figure S1.** FIB images of a BNNT-loaded muscle strip. The top images A-B) show the cuts made with the focused ion beam. C) Shows the longitudinal shape of a myotube in the matrix and D) a transversal one. E-F) show another myotube, where its membrane and a nucleus can be identified.

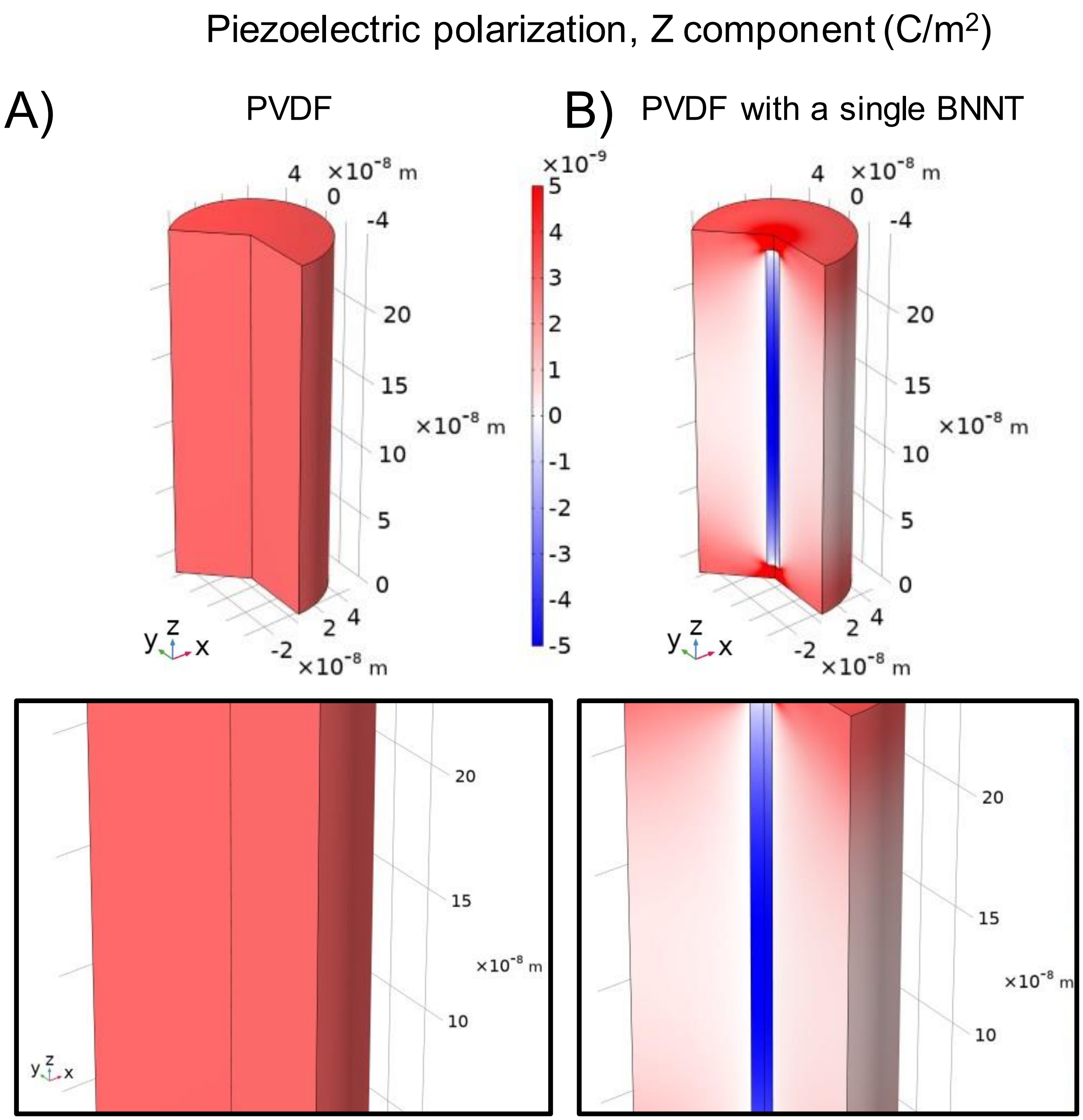


**Figure S2.** COMSOL simulation showing a single PVDF fiber (A) and a PVDF fiber with a BNNT inside (B). A compression effect of 100 Pa is applied along the z-axis and the piezoelectric polarization (C/m2) is represented. The piezoelectric polarization seen on the PVDF is positive while the one in the BNNT inside is negative, confirming the hypothesis we sustain about the inverse electric effect of both materials.

This supports our explanation on how the inverse polarization of both piezoelectric material respect to one another could be interfering in the dynamometer results. For this simulation, the d33 piezoelectric coefficient used for the BNNT was 0.76 C/m2 -as in the simulation of the biobot- and for the PVDF, -0.027 C/m2. The voltage produced by the PVDF fiber with a BNNT was of 6 µV while in the one without BNNT was 10 µV.

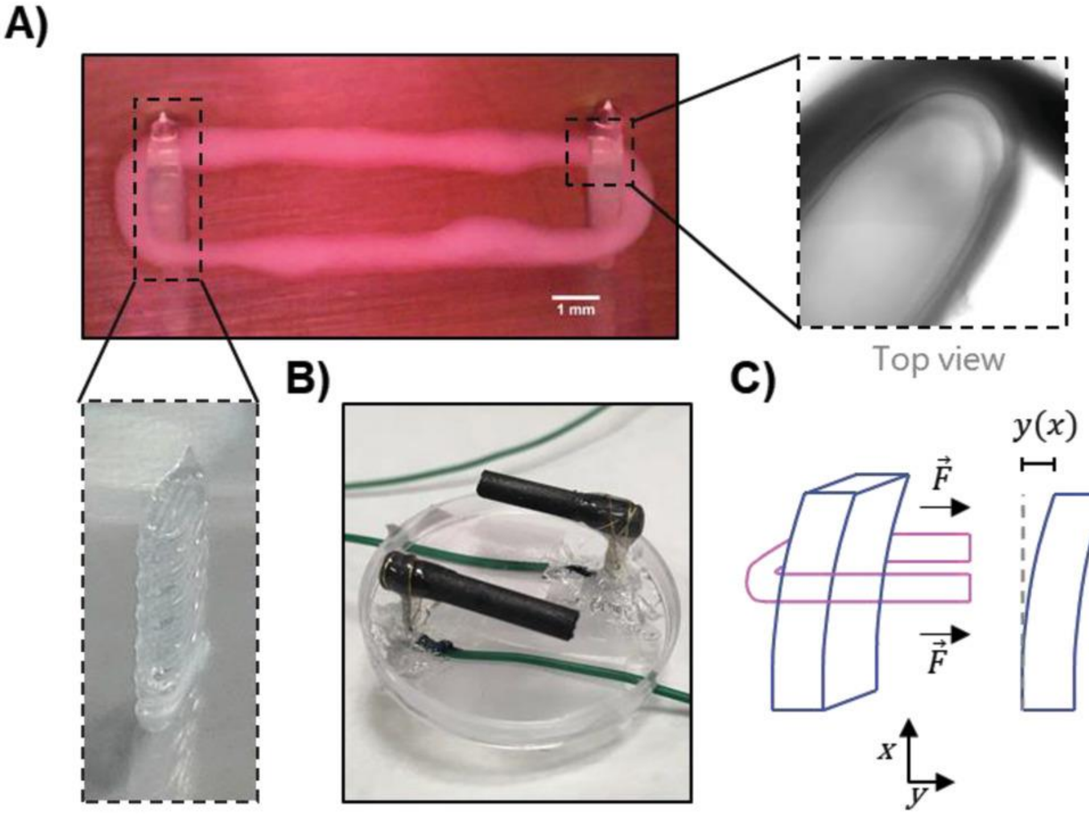


**Figure S3.** Two-post system used to measure the force of muscle tissues. A) The muscles were positioned surrounding two PDMS posts fabricated with 3D printing. Microscopic videos as in the to view were recorded by an inverted microscope when electric fields were being applied. B) A set of home-made carbon-based electrodes inserted into the lid of a Petri dish were used to electrically stimulate the samples. C) The force was calculated according to the displacement recorded in the microscope videos.

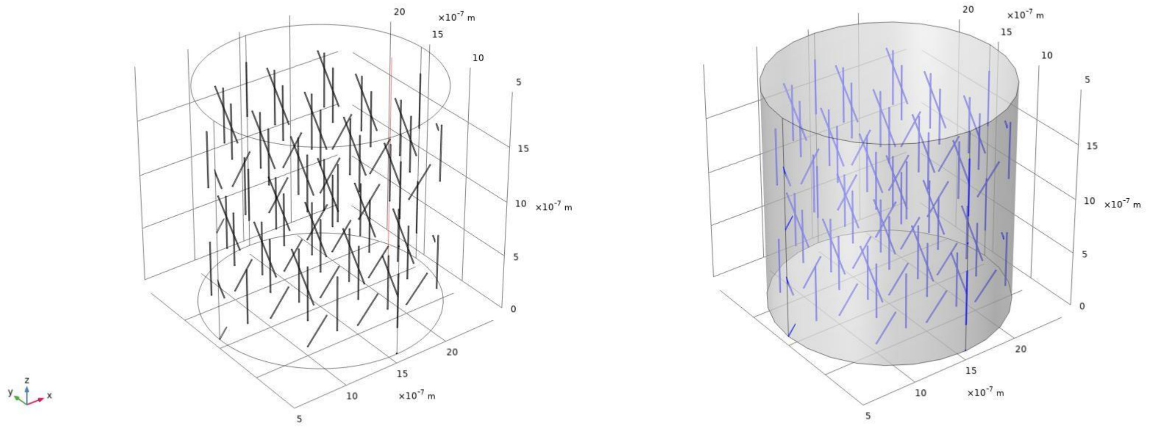

**Figure S4.** Images of the design of the simulation for the hydrogel-muscle matrix with embedded BNNTs in COMSOL. This design was performed reflecting a section of the biobot structure. The diameter of a single BNNT was established to be 10 nm, based on the SEM images obtained; and their length, 500 nm. In this simulation, all of the BNNTs have the same size but different orientations, representing their random disposition within the matrix among the muscle cells.

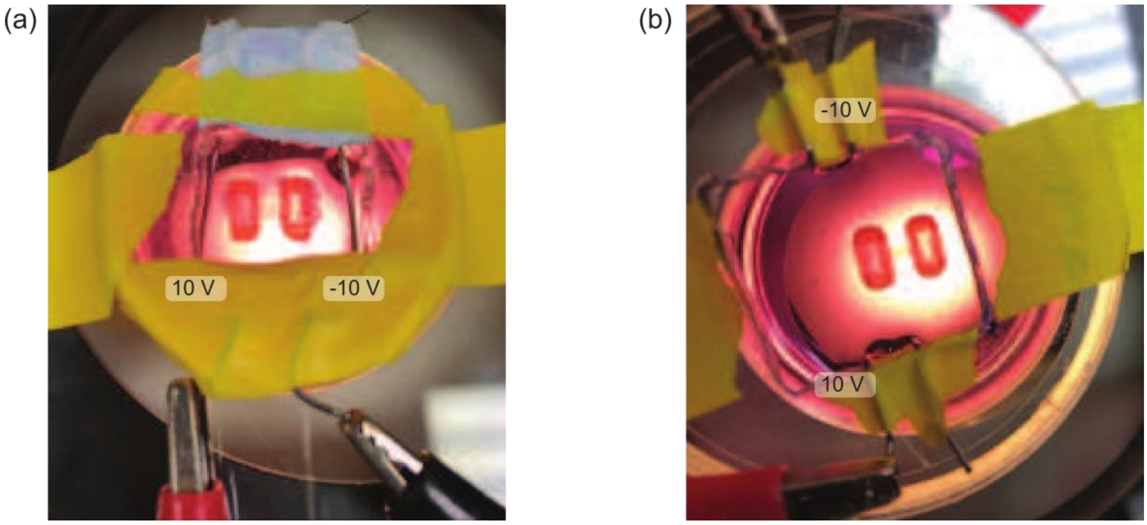


**Figure S5.** Stimulation setup showing the two platinum electrodes connected to the voltage generator and two bio-bots in the Petri dish.
